# Sex-specific resilience of neocortex to food restriction

**DOI:** 10.1101/2023.10.06.561185

**Authors:** Zahid Padamsey, Danai Katsanevaki, Patricia Maeso, Manuela Rizzi, Emily Osterweil, Nathalie L. Rochefort

## Abstract

Mammals have evolved sex-specific adaptations to reduce energy usage in times of food scarcity. These adaptations are well described for peripheral tissue, though much less is known about how the energy-expensive brain adapts to food restriction, and how such adaptations differ across the sexes. Here, we examined how food restriction impacts energy usage and function in the primary visual cortex (V1) of adult male and female mice. Molecular analysis and RNA sequencing in V1 revealed that in males, but not in females, food restriction significantly modulated canonical, energy-regulating pathways, including pathways associated with AMP-activated protein kinase (AMPK), peroxisome proliferator-activated receptor alpha (PPARα), mammalian target of rapamycin (mTOR), and oxidative phosphorylation. Moreover, we found that in contrast to males, food restriction in females did not significantly affect V1 ATP usage or visual coding precision (assessed by orientation selectivity). Decreased serum leptin is known to be necessary for triggering energy-saving changes in V1 during food restriction. Consistent with this, we found significantly decreased serum leptin in food-restricted males but no significant change in food-restricted females. Collectively, our findings demonstrate that cortical function and energy usage in female mice are more resilient to food restriction than in males. The neocortex, therefore, contributes to sex-specific, energy-saving adaptations in response to metabolic challenge.

## Introduction

Mammals reduce their energy usage in times of food scarcity. These adaptations have been well documented for peripheral tissues and are sex-specific. For example, females, as compared to males, readily lose muscle and bone mass to reduce peripheral energy expenditure during food restriction, but lose less fat mass and overall bodyweight^1–6^. Moreover, females are more likely to suppress energy-costly, reproductive functions during food restriction than males, evidenced by substantive reductions in uterine and ovarian mass, as well as a cessation of reproductive function and behaviour^7–9^. Such sex-specific differences are thought to reflect the differential importance of different organs to sex-specific survival^2^.

In contrast to peripheral tissue, whether and how the brain adapts its function and energy usage in a sex-specific manner during food scarcity remains largely unknown. The brain requires substantial amounts of energy to process and encode information^10–12^. Indeed, the human brain, which comprises 2% of our body’s mass, consumes 20% of our caloric intake, over half of which is used by the cerebral cortex^12^. Previously, we demonstrated that food restriction in male mice reduced energy usage in the primary visual cortex (V1) during visual processing, which was associated with a loss of orientation selectivity and visual coding precision^13^. These changes required a decrease in levels of serum leptin, a hormone secreted by adipocytes in proportion to fat mass^14^. Thus, in males the cortex, like peripheral tissue, has the capacity to reduce its energy usage and function in times of food scarcity.

By contrast, it remains unclear to what extent the brain similarly contributes to energy-saving adaptations during food scarcity in mammalian females. Understanding how food restriction impacts cortical function in males and females is not only of fundamental importance for understanding the sex-specific impact of diet on brain function, but is also critical for studies of cortical function, in which food restriction is used to behaviourally motivate animals^15–17^.

Here, we examined how food restriction leading to a 15% loss in body weight over 2-3 weeks impacts energy usage and visual coding in the primary visual cortex (V1) in adult male and female mice. We found that leptin levels, which regulate energy-saving changes in^13^, were maintained in females during food restriction, despite being decreased by approximately threefold in males. Consistent with these results, and using a range of experimental techniques, we failed to find significant energy-saving changes in V1 during food restriction in female mice, in contrast to male mice. Specifically, molecular analysis and RNA sequencing of V1 revealed that food-restriction significantly modified canonical energy-regulating cellular pathways in males, but not in females, including pathways associated with AMP-activated protein kinase (AMPK), peroxisome proliferator-activated receptor alpha (PPARα), and mammalian target of rapamycin (mTOR) signalling, as well as oxidative phosphorylation. Moreover, using *in vivo* ATP and calcium imaging, we found that food restriction significantly reduced V1 ATP usage and orientation selectivity in males, but not in females.

Collectively, our study demonstrates that in times of food restriction, visual cortical function and energy usage are largely maintained in female mice, while they are reduced in males. The neocortex, therefore, contributes to sex-specific, energy-saving adaptations in response to metabolic stress. Our research highlights the importance of taking into account sex differences when studying the impact of dietary manipulations on brain function.

## Results

We first examined sex-specific differences in how the neocortex adapts to food restriction in mice. Mice, 7-9 weeks of age, either had *ad libwitum* (CTR) access to food or were food restricted (FR) to 85% of their free feeding bodyweight, for 2-3 weeks (Figure 1A). Prior to any of our recordings, tissue collection, and analysis, all animals were given *ad libitum* access to food until sated. This enabled us to examine the long-term impact of caloric restriction on neocortical function, as opposed to the short-term impacts of hunger^18,19^.

**Figure 1.**
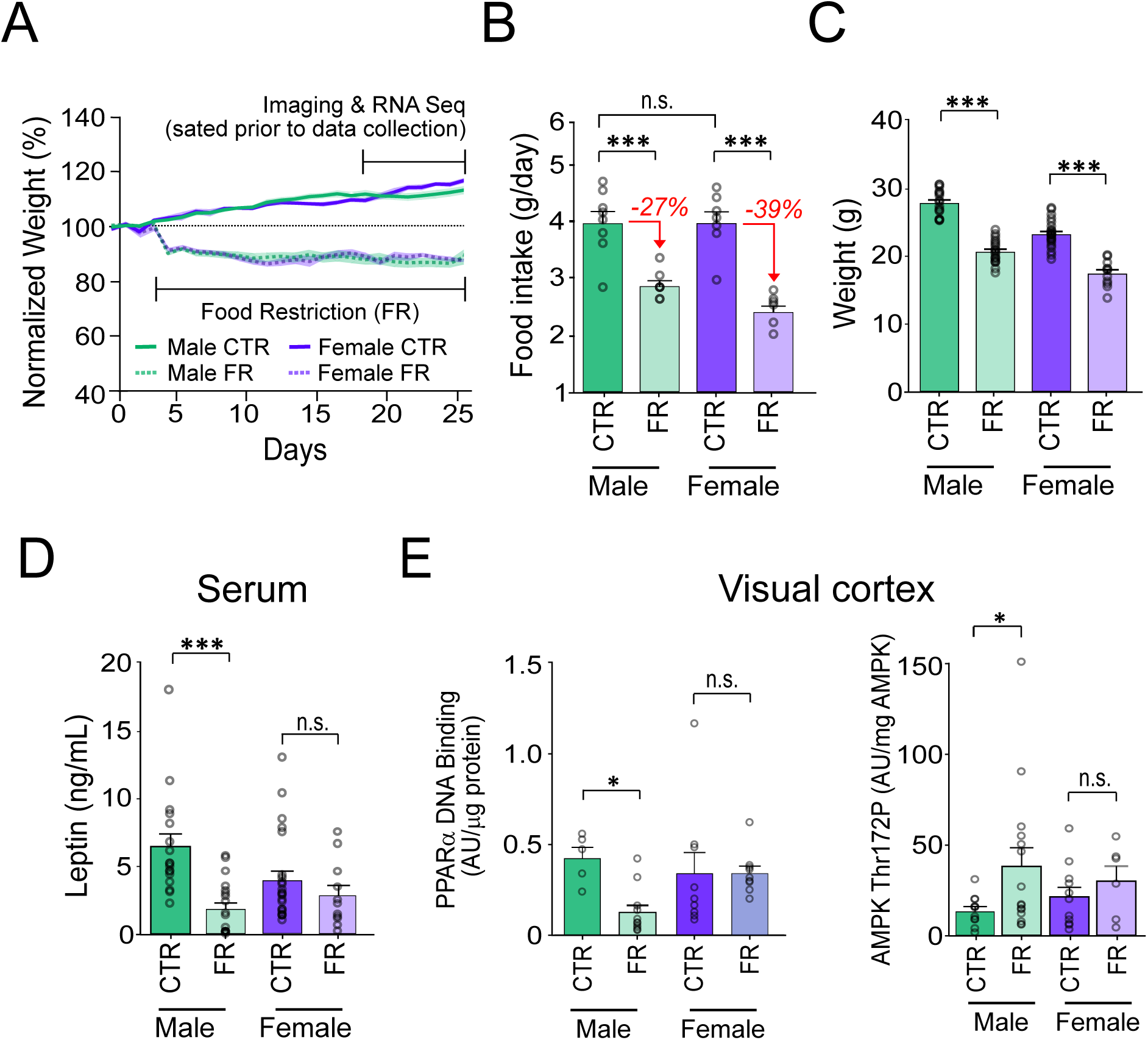
Sex-specific impact of food restriction on weight loss and serum leptin. (A) Animal weight across time. (B) Daily food intake (two-way ANOVA: CTR male vs FR male; t = 4.81; df = 25; p < 0.0001; CTR male vs. CTR female; t = 0.013; df = 25; p = 0.99; CTR female vs. FR female; t = 6.59; df=25; p < 0. 0001; FR male vs. FR female; t = 1.93; df = 25; p = 0.07; n = 8 CTR males, 8 FR males, 7 CTR females, and 7 FR females). Percent reduction of food intake for food restriction is shown in red for each sex. (C) Animal weight (two-way ANOVA: CTR male vs FR male; t = 11.36; df = 25; p < 0.0001; CTR female vs FR female; t = 8.36; df = 25; p < 0.0001; n = 17 CTR males and 19 FR males; 23 CTR females and 11 FR females). (D) Serum leptin levels (two-way ANOVA: CTR male vs FR male; t = 4.58; df = 66; p < 0.0001; CTR females vs FR females; t = 1.00; df = 66; p = 0.32; n = 17 CTR males and 19 FR males; 23 CTR females and 11 FR females). (E) Left. AMPK Thr172 phosphorylation, normalized by total AMPK, in primary visual cortex (V1) tissue (two-way ANOVA: CTR male vs FR male; t = 2.28; df = 39; p = 0.022; CTR female vs. FR female; t = 0.64; df = 39; p = 0.11; n = 11 CTR males, 15 FR males, 11 CTR females, and 6 FR females). Right. PPARα activity in V1 tissue, as assessed by levels of DNA binding, normalized to protein level (two-way ANOVA: CTR male vs FR male; t = 4.81; df = 30; p = 0.013; CTR female vs. FR female; t = 0.0016; df = 30; p = 0.99; n = 5 CTR males, 11 FR males, n = 9 CTR females, and 9 FR females). ***p < 0.0001; *p < 0.05; n.s. = not significant. Error bars are S.E.M.

### Sex-specific impact of food restriction on weight loss and fat mass-regulated hormone leptin

Prior to food restriction, we found that both males and females consumed the same *ad libitum* daily food intake, despite females weighing approximately 80% of age-matched males (weight: male vs. female; 27.82 g [95% CI: 26.84 g – 28.79 g] vs 23.21 g [95% CI: 22.35 g – 24.07 g]; t = 7.58; df = 66; p < 0.001; n = 8 male and 7 female mice). Critically, we found that to achieve a similar magnitude (15%) and rate of weight loss in both sexes we had to impose a 40% greater restriction of daily food intake in females (average of 39% of *ad libitum* intake; n = 7 mice) compared to males (27% of *ad libitum* intake; n = 8 mice) (Figure 1B,C). This is consistent with previous studies demonstrating that female mice are more resistant to weight loss than male^1–6^. Moreover, for similar bodyweight loss (15%) maintained over 2-3 weeks, serum leptin levels, which reflect fat mass, were markedly and significantly decreased (-72%) in males (leptin: CTR male vs FR male; 6.45 ng/mL [95% CI: 4.47 ng/mL – 8.43 ng/mL] vs. 1.84 ng/mL [95% CI: 0.90 ng/mL – 2.79 ng/mL]; t = 4.58; df = 66; p < 0.0001; n = 17

CTR males and 19 FR males), but only modestly and non-significantly decreased (-28%) in females (leptin: CTR females vs FR females; 3.94 ng/mL [95% CI: 2.53 ng/mL – 5.35 ng/mL] vs. 2.83 ng/mL [95% CI: 1.20 ng/mL – 4.47 ng/mL]; t = 1.00; df = 66; p = 0.32; 23 CTR females and 11 FR females) (Figure 1D). This is in keeping with previous findings that female mice are more resistant to fat loss than males^1–6^ and that weight loss arises from other tissues including those associated with reproductive functions (ovarian and uterine mass)^7–9^.

### Cellular energy-regulating pathways in cortex are significantly impacted by food restriction in males but not in females

Decreased leptin levels are necessary for triggering energy-saving changes in the mouse visual cortex during food restriction^13^. We therefore asked whether the absence of significant change in serum leptin in food-restricted females was associated with any change in V1 energy usage. We first assessed this at the molecular level by examining AMPK phosphorylation state. AMPK is a canonical cellular energy sensor activated by metabolic stress - such as food restriction, in part via decreased leptin signalling - through phosphorylation of threonine 172 (Thr^172^)^20–22^. Its activation results in reduced ATP expenditure, for example, through the inhibition of mammalian target of rapamycin (mTOR) signalling^20,23^, and regulation of mitochondrial oxidative phosphorylation^24,25^. In V1, we found that AMPK Thr^172^ phosphorylation was markedly and significantly elevated (2.9 fold) with food restriction in males (AMPK Thr^172^ phosphorylation: CTR male vs FR male; 13.19 AU/μg [95% CI: 7.68 AU/μg – 18.71 AU/μg] vs. 38.21 AU/μg [95% CI: 16.50 AU/μg – 59.93 AU/μg]; t = 2.28; df = 39; p = 0.022; n = 11 CTR males and 15 FR males), consistent with ATP savings, but only modestly increased (1.4 fold) in females; this increase was not statistically significant (AMPK Thr^172^ phosphorylation: CTR female vs. FR female; 20.94 AU/μg [95% CI: 9.42 AU/μg - 32.46 AU/μg] vs. 29.52 AU/μg [95% CI: 8.52 AU/μg – 50. 53 AU/μg]; t = 0.64; df = 39; p = 0.11; 11 CTR females, and 6 FR females) (Figure 1E). We also examined the activity of peroxisome proliferator-activated receptor alpha (PPARα), which is downstream of leptin signalling and is known to regulate fatty acid metabolism and energy homeostasis^22,26^. As with AMPK, we found that PPARα was significantly regulated by food restriction selectively in males, but not in females (Figure 1E)^27,28^.

To further investigate how energy-regulating cellular pathways are impacted by food restriction, we performed RNA sequencing on tissue dissected from V1 (Figure 2A; n = 4 CTR male, n = 4 FR male, n = 4 CTR female, and n = 4 FR female mice). Differential gene expression analysis using DESeq2 revealed that food restriction significantly altered the expression of 657 and 585 targets in males and females, respectively (at a threshold of p-adj ≤ 0.1; only 125 of these were common to both sexes (Figure 2B). To assess the functional relevance of the transcriptomic changes induced by food restriction, we performed Gene Set Enrichment Analysis (GSEA) to identify pathways enriched in the differentially expressed populations. We found that food restriction resulted in a significant alteration of 34 and 14 gene sets (at p-adj ≤ 0.05) in males and females respectively; 13 of these were common to both sexes (Figure 2C; Table 1; Table 2). In male mice, we specifically found that food restriction significantly regulated gene sets associated with canonical energy-regulating pathways (Figure 2), including those associated with oxidative phosphorylation and mTOR signalling (mTORC1 signalling and PI3K/AKT/mTOR signalling), as well as fatty acid metabolism; these pathways are known to regulated by leptin and AMPK signaling^20,27–29^ (Figure 2D). Notably, these pathways were not significantly regulated by food-restriction in females. In addition to these findings, we found that gene sets associated to estrogen-regulating pathways were significantly regulated in food-restricted males and not in females (Figure 2C). This is consistent with previous findings showing that estrogen, like leptin, suppresses food intake and promotes energy expenditure^30^. Finally, consistent with past studies in cortex, we found that food restriction in males altered gene expression associated with inflammation and reactive oxidative species (Figure 2D)^31,32^.

**Figure 2.**
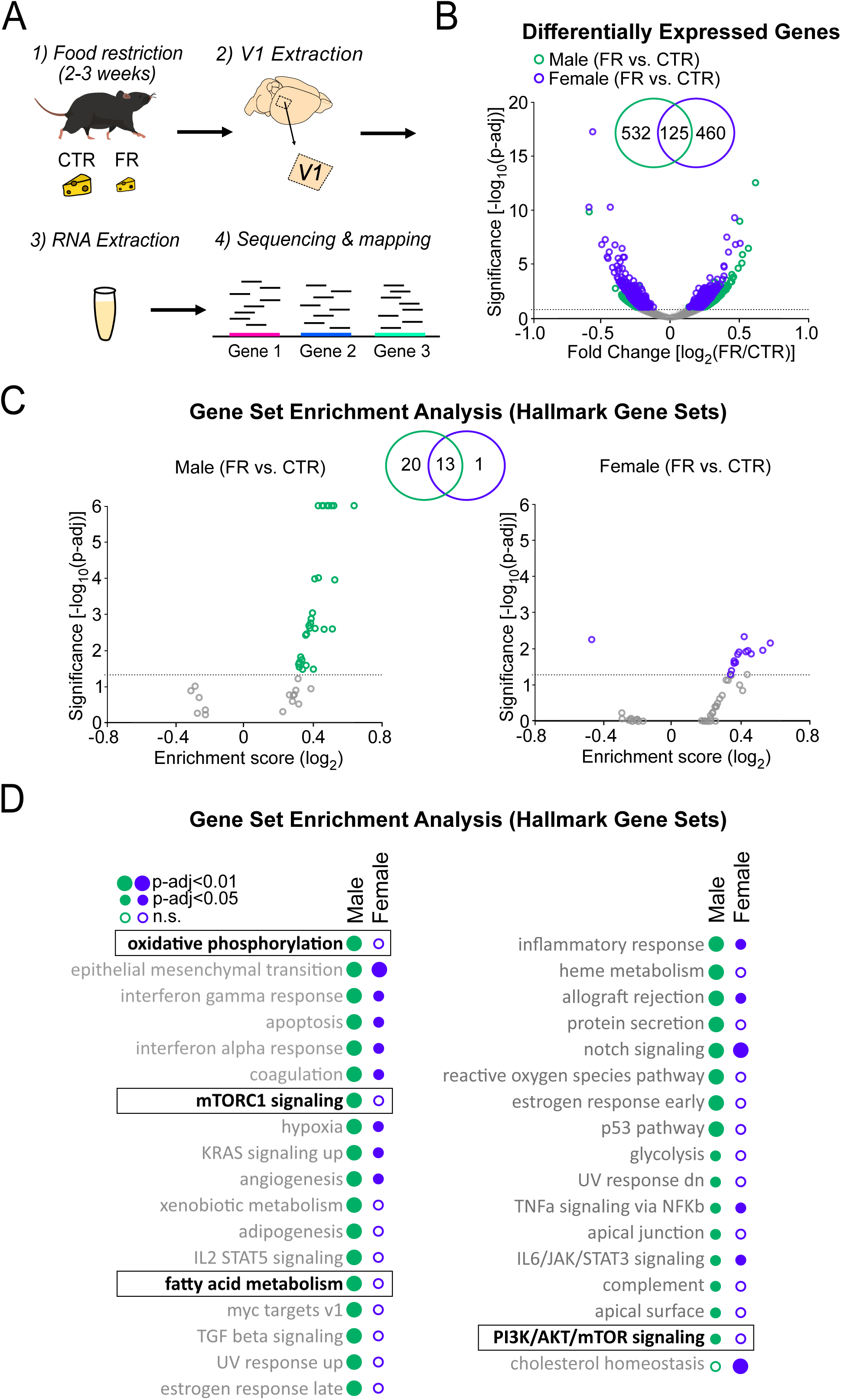
Cellular energy-regulating pathways are more robustly impacted by food restriction in males than in females. (A) Schema of RNA sequencing of V1 tissue. (B) Volcano plot of fold change expression with food restriction vs. significance for all analyzed genes; broken horizontal line denotes adjusted p ≤ 0.1 significance level, below which non-significantly regulated expressed genes are marked by grey. Inset: Venn diagram showing the number of significantly differentially expressed genes with food restriction in males and females. (C) Gene Set Enrichment Analysis using Hallmark Gene Sets, depicted using a volcano plot of Enrichment Score vs. significance for males (left) and females (right); broken horizontal lines denotes adjusted p < 0.05 significance level. Inset: Venn diagram showing the number of significantly regulated gene sets with food restriction in males and females. (D) Complete list of significantly enriched gene sets by food-restriction in males and females for analysis in C. Gene sets of interest are denoted by a box (oxidative phosphorylation: male; t = 0.48; p < 0.0001; female; t = -0.17; p = 0.96; mTORC1 signaling: male; t = 0.46; p < 0.0001; female; t = -0.18; p = 0.98; fatty acid metabolism: male; t = 0.43; p < 0.0001; t = 0.2; p = 0.88; PI3K/AKT/mTOR signaling: male; t = 0.34; p = 0.027; female; t = 0.17; p = 0.98). All data from 4 CTR males, 4 FR males, 4 CTR females, and 4 FR females.

Collectively these findings reveal that food restriction strongly affects energy-regulating cellular pathways in the cortex of male, as opposed to female, mice.

### ATP usage in V1 is significantly decreased under food restriction in males but not in females

We next examined energy usage in visual cortex during visual processing using two photon ATP imaging in V1 of awake, head-fixed Thy1-ATeam1.03^YEMK^ mice, which express a FRET-based ATP sensor^33,34^ (Figure 3A). In the presence of ATP synthesis inhibitors, focally applied over the visual cortex, ATP usage can be seen as a decay of the FRET signal during visual stimulation with natural stimuli^13^ (Figure 3A). Data obtained in females were compared to our previously published datasets obtained under similar conditions in male mice^13^. As previously shown^13^, food restriction robustly decreased the rate of cortical ATP usage in males (24% decrease) (half-time of FRET decay: CTR male vs. FR male; 8.48 min [95% CI: 7.59 min – 9.37 min] vs. 10.51 min [95% CI: 9.08 min – 11.93 min]; t = 2.87; df = 37; p = 0. 0067; n = 11 CTR males and 10 FR males) during visual stimulation; a more modest trend was observed in females (12% decrease), which was not significant compared to controls (half-time of FRET decay: CTR female vs. FR female; 8.66 min [95% CI: 7.49 min – 9.84 min] vs. 9.66 min [95% CI: 8.92 min – 10.39 min]; t = 1.36; df = 37; p = 0.18; n = 12 CTR females and n = 8 FR females) (Figure 3B-C). In the absence of visual stimulation (darkness), ATP usage was lower than during visual stimulation and unaffected by food restriction, for either sex (Supplemental Figure 1).

**Figure 3.**
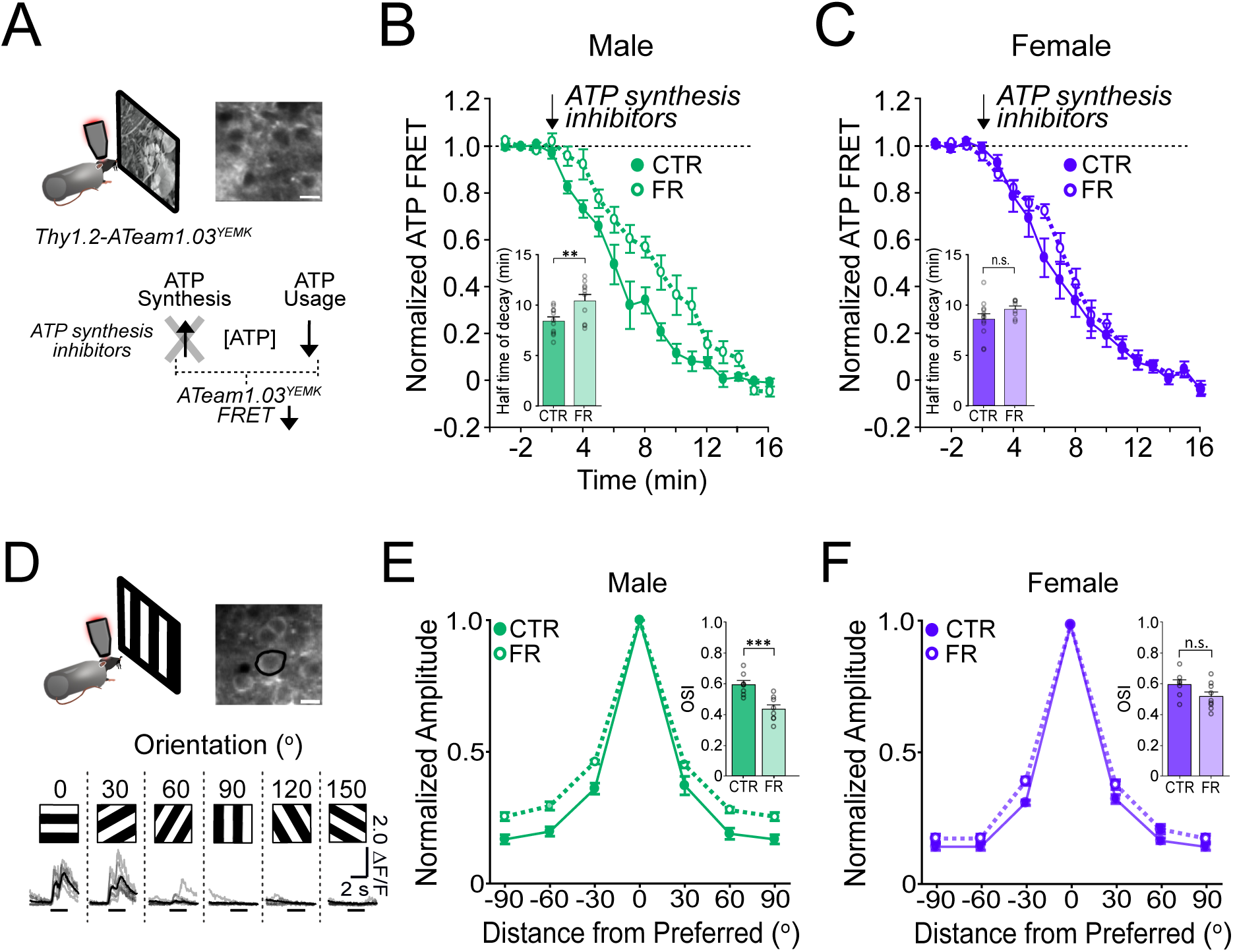
ATP usage and orientation selectivity in V1 are more robustly decreased by food restriction in males than in females. (A) Schemata of ATP imaging experiment. Top: example field of view of V1 layer 2/3 neurons in the ATeam1.03^YEMK^ transgenic mouse (scale bar: 10 μm). Bottom: ATP synthesis inhibitors were used to isolate ATP usage, recorded as a decrease in FRET signal during presentation of natural, outdoor scenes. (B) Normalized ATeam1.03^YEMK^ FRET signal in males and during presentation of natural, outdoor scenes. ATP synthesis inhibitors (arrow) were added to isolate ATP usage. Inset: Time to 50% decay of the ATeam1.03^YEMK^ FRET signal (two-way ANOVA: t = 2.87; df = 37; p = 0. 0067; n = 11 CTR males and 10 FR males). (C) As in (B) but for females (inset: two-way ANOVA: t = 1.36; df = 37; p = 0.18; n = 12 CTR females and n = 8 FR females). (D) Top: Schema of two-photon imaging and sample field of view of V1 layer 2/3 neurons expressing GCaMP6s (scale bar: 10 μm). Bottom: Sample fluorescent signals (grey) from a selected neuron (black circle) in response to 6 drifting gratings of varying orientations. Trial averages are in black. Horizontal bar denotes 2s grating presentation. (E) Mean orientation tuning curves normalized to the response to the preferred orientation for CTR and FR males (data from ^13^). Note that -90° and +90° conditions correspond to the same visual stimulus. Inset: Orientation selectivity index (OSI; two-way ANOVA: t = 3.75; df = 27; p = 0.0009 n = 8 CTR and 8 FR males). (F) As in (E) but for females (Inset: two-way ANOVA: t = 1.90; df = 27; p = 0.08; n = 7 CTR females and 9 FR females). **p < 0.01; ***p < 0.001; n.s. = not significant. Error bars are S.E.M. Data from male mice are from a previously published dataset obtained under similar conditions^13^.

Altogether, these findings demonstrate that food-restriction significantly impacts V1 energy usage during visual stimulation in males but not in females.

### Orientation selectivity in V1 is significantly reduced under food restriction in males but not in females

We next examined how food restriction impacts visual cortical function. In male mice, reductions in energy usage in V1 during food restriction are associated with a decreased visual coding precision as evidenced by a reduction of orientation selectivity^13^. Given the non-significant impact of food-restriction on energy usage in female mice, we reasoned that orientation selectivity would also be less impacted. We analysed two-photon calcium imaging data of V1 GCaMP6s-labeled layer 2/3 neurons in female mice viewing drifting gratings (Figure 3D-F; n = 7 CTR male, n = 8 FR male, n = 7 CTR female, and n = 9 FR female mice). Data obtained in females were compared to our previously published datasets obtained under similar conditions in male mice^13^. Consistent with our hypothesis, in contrast to males in which orientation selectivity was significantly reduced by 27% (OSI: CTR male vs. FR male; 0.58 [95% CI: 0.51 – 0.65] vs. 0.42 [95% CI: 0.36 – 0.49]; t = 3.75; df = 27; p = 0.0009), orientation selectivity was modestly (13%) and not significantly reduced by food-restriction in females (OSI: CTR female vs. FR female; 0.60 [95% CI: 0.52 – 0.67] vs. 0.52 [95% CI: 0.45 – 0.58]; t = 1.90; df = 27; p = 0.08) (Figure 3E-F). Direction tuning, as measured by the direction selectivity index (DSI) was also significantly affected by food-restriction in males, but not in females (Supplemental Figure 3; Males: CTR vs. FR; t = 2.16; p = 0.023; df = 12; n = 6 CTR males and 8 FR males; Females: CTR vs. FR; t = 1.49; p = 0.16; df = 14; n = 7 CTR females and 9 CTR females).

Altogether, our results show that energy usage and coding precision in the visual cortex are largely maintained in female mice under food restriction while they are robustly reduced in males. Such findings cannot be readily attributed to differences in behavioral or attentional state. All animals were imaged whilst in a cardboard tube, and therefore not locomoting. Moreover, we found no difference in pupil size during the presentation of visual stimuli (Supplemental Figure 2).

### Bayes factor analysis reveals statistically robust impact of metabolic stress on V1 energy usage and visual coding in males, but not females

We used Bayes factor hypothesis testing to better assess the statistical impact of food restriction on visual cortical function and energy usage in male and female mice (Supplemental Figure 3). Bayes factor analysis uses a Bayesian framework to quantify how much more likely the data is under the alternative hypothesis (*i.e.* there is an effect of food restriction) versus the null hypothesis (*i.e.* there is no effect of food restriction), quantified by the Bayes factor (BF_10_)^35,36^. A BF_10_ > 3, for example, means that the data are >3 times more likely under the alternative hypothesis than the null hypothesis. Critically, the BF_10_ allows for a more informative comparison of the effects of food restriction between sexes, as opposed to comparing p-values, which can be misleading^35,36^. Across the parameters of energy usage and visual coding that we tested (Figures 1-3), BF_10_ for males was consistently several-fold greater than that for females. Thus, visual cortical function and energy usage was more robustly modulated by food restriction in males than in females.

Our findings however cannot rule out a potential, smaller impact of food restriction on cortical function and energy usage in females, below the threshold for statistical detection in this study. In particular, for females, we found only 3 out of the 10 parameters had BF_10_ < 0.33, *i.e.* the data are 3 times more likely under the null hypothesis than the alternative hypothesis supporting the absence of an effect ^35,36^(Supplemental Figure 3). This suggests that food restriction may have a modest impact on V1 energy usage or function in females, but below the limits of statistical detection in our study.

Altogether, our results show that energy usage and coding precision in the visual cortex are largely maintained in female mice under food restriction while they are robustly reduced in males.

## Discussion

Here, we find that in female mice, energy usage and visual coding in V1 is largely unchanged under food restriction. Using molecular analysis and RNA sequencing, we found that energy-regulating pathways were not significantly changed with food restriction in females, in contrast to males; these included pathways associated with AMPK, mTOR, and PPARα signalling, as well as oxidative phosphorylation. Consistent with this, we found that food restriction in females did not impact visual cortical ATP usage nor orientation selectivity, both of which were reduced by food restriction in males. Our findings suggest that the contribution of neocortex to energy-saving adaptations during food restriction is sex-selective and more pronounced in males; females may rely on other energy-saving strategies, such as reducing reproductive function, to cope with food restriction.

### Limitations of the study

Our study focused on the effects of a moderate food restriction protocol over 2-3 weeks on neocortical function and energy usage. This protocol is widely used in metabolic studies investigating the impact of caloric restriction on peripheral physiology, as well as in neuroscience studies to motivate rodents to perform specific behaviors^6,15–17^. However, it remains unclear to what extent our findings generalize to other protocols of food restriction. Indeed, the duration, magnitude, and pattern of food restriction, as well as macronutrient composition are known to impact the physiological responses to dietary manipulations^22,37–39^. Moreover, whilst we focussed our study on the adult visual cortex, it remains unclear if the brain shows similar sex-specific energy-saving adaptations in other cortical areas, and how such adaptations change with age^6,40^.

A key strength of our study is that we consistently found, across a number of experimental methods, that changes in cortical function and energy usage were significant in males, but not in females. However, a critical limitation of our study is that we did not have the statistical power to directly compare the effect size of food restriction between the sexes (e.g. ANOVA interaction effects); to do so would require a substantial increase in experimental group sizes by several fold. We addressed this limitation, in part, using Bayes factor analysis, which revealed that the impact of food restriction in males was indeed more robust than in females. This analysis also revealed that our study did not have sufficiently strong evidence to definitively rule out a smaller effect of food restriction in females, which may not have been statistically detectable.

### Role of leptin and sex hormones in food restriction impact on cortical activity

Consistent with past studies, we found that food-restricted female mice resisted weight loss, and therefore required a greater restriction of food to achieve similar weight loss (relative to baseline) as males. In addition, for the same relative weight loss as males, females exhibited minimal reductions in serum leptin levels (secreted by adipocytes), consistent with past studies showing that females resist fat loss during food restriction^1–3,5,6^. Similar effects have been observed across several mammalian species, including humans, and is thought to reflect the differential importance of fat stores for reproductive success in males and females^1–5^. Notably, we previously demonstrated that reductions in leptin levels during food restriction are necessary for driving energy-saving changes in visual cortical coding in male mice; leptin therefore acts as a critical signal linking the neocortex to fat stores^13^. That leptin levels are not significantly affected by food restriction in females may explain the minimal impact food restriction has on cortical energy usage and coding precision in female mice.

In addition to leptin, sex hormones likely contribute to sex-specific adaptations to food restriction^41^. As with leptin, estrogen suppresses food intake and promotes energy expenditure^30^, in part by augmenting leptin signalling by increasing leptin-induced phosphorylation of STAT3^42^. Moreover, estrogen can enhance ATP production by sensitizing insulin signalling, enhancing glycolysis and oxidative phosphorylation, and along with progesterone, upregulating mitochondrial gene expression and function^30,43,44^. Interestingly, in our study, RNA sequencing revealed that estrogen signalling pathways were significantly regulated in male visual cortex with food-restriction, and were unchanged in females. Estrogen may thus have overlapping effects with those induced by food restriction, thereby diminishing further responses to food restriction in females^6^. Notably, it was recently shown that sex differences in calorie-restriction’s metabolic effects that are observed in young adult mice, are largely absent in older mice, when females’ estrogen levels have declined^6^. Further studies will be needed to explore whether the resilience of cortical function found in this study in young cycling adult females is also observed in non-cycling mice with reduced oestrogen levels, such as in aged mice.

### Sex-specific regulation of cellular energy-regulating pathways

AMPK and mTOR signalling pathways are canonical regulators of cellular energy usage^20,23^. We found these pathways, along with cortical ATP usage, to be significantly modulated by food-restriction in male, but not female mice. AMPK is activated during metabolic stress, and saves energy by regulating oxidative phosphorylation^24,25^, and inhibiting cellular pathways associated with energy usage, such as mTOR signalling^20^. mTOR signalling, by upregulating protein synthesis, increases ATP usage by promoting neuronal excitability, neurite outgrowth, ion channel expression, and synaptic plasticity^23^. mTOR signalling is therefore ideally situated to regulate how much ATP is used in neuronal function; its inhibition may underlie ATP savings and the associated loss of coding precision with food-restriction in male mice. Notably, previous studies have found sex- and tissue- dependent regulation of AMPK and mTOR signalling, including in response to food restriction^41^. For example, in skeletal muscle, mTOR signalling is enhanced by fasting selectively in male, but not female mice^41,45^. Moreover, metabolic challenge by endurance training induces greater AMPK activity in cardiac tissue in male, as opposed to female mice^46^. Our study extends the sex-specific regulation of these energy-regulating pathways to the neocortex.

### Importance of biological sex on the impact of food restriction on cortical function

Food restriction protocols are widely used to motivate animals in behavioural studies examining cortical function^15–17^. Our findings suggest that these protocols are likely to have greater impacts on basal neocortical function in males, as compared to females. Consistent with this, a recent study has found that food restriction in juvenile mice strongly impacts cognitive flexibility in learning and decision-making in adulthood, specifically for males but not females^47,48^. Such sex-specific impacts of food restriction on neocortical function are important to take into consideration in the design and interpretation of such studies.

Diet is known to have a considerable impact on neurological diseases, including epilepsy, Alzheimer’s disease, and other neurodegenerative diseases^49,50^. Whilst calorie restricted diets have proven to be of benefit in this regard, the sex-dependency of such interventions have not been thoroughly investigated, though on the basis of our findings it is conceivable that males and females may respond differently to dietary interventions. The inclusion of both males and females in biomedical research, and the study of sex differences, is therefore of considerable importance for ensuring efficacious treatments are developed for both sexes.

In conclusion, we find that visual cortical coding and energy usage in females are resistant to food restriction, in contrast to males. Our findings establish that sex-specific adaptations in energy usage, previously described in peripheral tissue under metabolic challenge, extends to the neocortex.

## Acknowledgments

We thank the GENIE Program and the Janelia Research Campus (V. Jayaraman, R. Kerr, D. Kim, L. Looger, and K. Svoboda) for making GCaMP6 available. We thank J. Hirrlinger (Carl Ludwig Institute of Physiology, University of Leipzig, Germany) for providing the ATeam1.03^YEMK^ mouse line. We thank Sang Seo and Aditi Singh for assistance and guidance in RNA sequencing and analysis. We thank Will Cawthorn for discussions on the manuscript. This work was funded by the BBSRC (Responsive Mode Research Grant (BB/T007907/1 to N.R. and Z.P.), the Royal Commission for the Exhibition of 1851 (research fellowship to Z.P.), the MRC (G116854; Career Development Fellowship to Z.P.), the Wellcome Trust and the Royal Society (Sir Henry Dale fellowship to N.R.), the Shirley Foundation, the Patrick Wild Centre, the RS MacDonald Charitable Trust Seedcorn Grants (to N.R. and Z.P.), the Simons Initiative for the Developing Brain (to N.R. and D.K.) and the Wellcome Trust-University of Edinburgh Institutional Strategic Support Fund (ISSF3) (to N.R.). This project has received funding from the European Research Council (ERC) under the European Union’s Horizon 2020 research and innovation programme (grant agreement No. 866386).

## Author contributions

Z.P. and N.R. designed the experiments. Z.P. performed and analysed calcium imaging experiments and ELISAs. D.K. performed and analysed calcium and ATP imaging experiments. Z.P. and M.R. and E.O. processed tissue for RNA sequencing and analyzed data. P.M. performed and analyzed ELISAs. N.R. supervised the project. Z.P. and N.R. wrote the manuscript with input from all authors.

## Declaration of Interests

The authors declare no competing interests.

## Inclusion and Diversity

One or more of the authors of this paper self-identifies as an underrepresented ethnic minority in science. One or more of the authors of this paper self-identifies as a member of the LGBTQ+ community. While citing references scientifically relevant for this work, we also actively worked to promote gender balance in our reference list.

## STAR METHODS

### RESOURCE AVAILABILITY

#### Lead contact

Further information and requests for resources and materials should be directed to and will be fulfilled by the lead contact Nathalie L. Rochefort.

#### Materials availability

This study did not generate new unique reagents.

#### Data and code availability

1. GEO accession number for RNA-seq datasets: GSE233435
2. Processed data will be made available at https://datashare.ed.ac.uk/handle/10283/3871. Requests for raw data should be made to and will be fulfilled by the lead contact.
3. MATLAB scripts to analyse data have been previously published^13^ and are available at https://zenodo.org/record/5561795#.YmJ3AtrMJPY and https://github.com/rochefort-lab/Padamsey-et-al-Neuron-2022.
4. Any additional information required to reanalyze the data reported in this paper is available from the lead contact upon request.

## EXPERIMENTAL MODEL AND SUBJECT DETAILS

Experiments were approved by the University of Edinburgh’s Animal Welfare and Ethical Review Board (AWERB) and carried out under Home Office (UK) approved project and personal licenses. All experiments conformed to the UK Animals (Scientific Procedures) Act 1986 and the European Directive 86/609/EEC and 2010/63/EU on the protection of animals used for experimental purposes.

This study used male and female C57BL/6J mice (RRID:IMSR_JAX:000664; Jackson Laboratory) and male B6-Tg (Thy1.2-ATeam1.03^YEMK^)^AJhi^ transgenic mice, that were bred on a C57BL/6J background (RRID:MGI:5882597; https://scicrunch.org/resources). Animals were group housed (2-5 animals/cage) and maintained on a reverse light/dark (12h/12h) cycle in a room kept at 21±2°C and 55±10% humidity.

For data associated with *in vivo* recordings (Figure 3), data obtained in female mice were compared to our previously published datasets obtained under similar conditions in male mice^13^

## METHOD DETAILS

### Food restriction

Mice (7-9 weeks of age) either had *ad libitum* access to food (RM1 expanded pellets; DBM Scotland UK) or were food restricted for 2-3 weeks to 85% of their baseline bodyweight prior to experimentation as previously described^13^. Briefly, for food restriction, one ration of food was given 4-8 hours prior to the end of the dark cycle, with the amount of food progressively reduced to achieve target weight. 1-2 hour prior to experimentation, all animals had *ad libitum* access to food until sated.

### AAV injection and cranial window

For calcium imaging experiments, C57BL6 were anesthetized using isoflurane, and given pre-operative analgesia (vetergesic: 0.1 mg/kg; carprieve: 5 mg/kg; rapidexon: 2 mg/kg) along with warm Ringer’s solution (25ml/kg) subcutaneously; vetergesic jelly was additionally given orally post-operatively 24 hours later. Opaque eye cream was applied to the eyes (Bepanthen, Bayer, Germany). A craniotomy (2x2 mm) was applied over left V1 (centre at 2.5 mm mediolateral and 0.5 mm anterior to lambda). Flexed GCaMP6S (AAV1.Syn.Flex.GCaMP6s.WPRE.SV40; Addgene; RRID:Addgene_105558-AAV1) diluted 1:10 in saline, along with CaMKII-dependent Cre-recombinase (AAV1.CamKII 0.4.Cre.SV40; Addgene; RRID:Addgene_105558-AAV1) diluted 1:100 saline was injected at the centre of the craniotomy via a sharp glass pipette. Injections were targeted to layer 2/3, with 100 nL of virus solution injected (2.5nL/30s) at each of 3 depths (150, 250 and 350 μm) via a sharp glass pipette using a Nanoject III (Drummond Scientific). A glass window was then used to seal the craniotomy, and superglued in place. The window was made of 1 or 2 glass coverslips (Menzel-Glaser # 0), in the latter case the 2 cover slips were glued together with optically-clear, UV-cured glue (Norland Optical Adhesive no. 60); the inner window was 2.0 mm x 2.0 mm, the outer window was 2.5 mm x 2.5 mm. A metal headplate was then fixed to the skull with super glue and dental cement (Paladur, Heraeus Kuzler). It took 3-6 weeks for viral expression prior to experimentation.

For ATP imaging experiments, we used ATeam1.03^YEMK^ mice (RRID:MGI:5882597; https://scicrunch.org/resources) that express a FRET-based sensor under the Thy1 promoter^33^. One the day of experimentation, animals were anesthetized with isoflurane and given pre-operative analgesia (Carprieve: 5 mg/kg) and 25 mL/kg of warm Ringer’s solution subcutaneously. A headplate was first fixed to the skull with superglue and dental cement, after which a small craniotomy (∼0.5 x ∼0.5 mm) was made over left V1 (centre at 2.5 mm mediolateral and 0.5 mm anterior to lambda), which was covered with agarose (1-2%) and silicone. Following recovery from anaesthesia (30-60 minutes) the animal was head-fixed; the agarose and silicone were removed and replaced with HEPES-buffered ACSF (in mM: 124 NaCl, 20 Glucose, 10 HEPES, 2.5 KCl, 1.2 NaH_2_PO_4_, 2 CaCl_2_, and 1 CaCl_2_; pH 7.2-7.4).

### Habituation and Head-fixation

During imaging, mice were placed in a cardboard tube and head fixed. Prior to the imaging day, mice were handled daily for 2-3 weeks prior, and exposed to cardboard tubes, similar to the one on the experimental rig. They were also habituated to the rig via once daily, 10-15 minute sessions for 2 days prior to recording. For ATP imaging, mice were habituated to the experimental setup without head fixation (10-15 min session/day for 1-2 days).

### *In vivo* two-photon imaging

ATP and calcium imaging were using a two-photon resonant scanning microscope, as previously described^51–53^, which was equipped with a Ti:Sapphire laser (Charmeleon Vision-S, Coherent, CA) and GaAsP photomultiplier tubes (Scientifica), and controlled with LabView (v8.2; National Instruments, UK). Time-series xy images were acquired at 40Hz using a 25x water-immersion objective (Nikon; CF175 Apo 25XC W; 1.1 NA) at depths between 160 and 280 μm below the pia. For GCaMP6s imaging, excitation was tuned to 920nm. For ATP imaging (CFP/YFP FRET sensor), the laser was tuned to 850 nm for CFP excitation. CFP and YFP fluorescence were recorded simultaneously using a 515nm long-pass dichroic mirror with the following emission filters: 485/70nm for CFP and 535/45nm for YFP (mirror and filter set: T515lpxr C156624; Scientifica). Imaging data were acquired during the presentation of visual stimuli, with at least 10 replicate trials for each visual stimulus. For ATP imaging, three trials were taken at baseline, then ACSF containing ATP synthesis inhibitors (1 mM oligomycin and 20 mM sodium iodoacetate) was applied over the open craniotomy isolate ATP usage. Thirty trials were successively performed immediately after drug application. For in vivo recordings, data obtained in females were compared to our previously published datasets obtained under similar conditions in male mice^13^.

### Visual Stimuli

MATLAB (Mathworks; RRID:SCR_001622; Psychophysics Toolbox; RRID:SCR_002881) was used to generate drifting grating stimuli. These were displayed on an LCD monitor (51 x 29 cm; Dell) placed 20 cm from the right eye, contralateral to the hemisphere with the cranial window. For food restriction experiments, on a given trial, 12 full-field drifting gratings (drift angle: 0, 30, 60, 90, 120, 180, 210, 240, 270, 300, 330°; temporal frequency = 1 Hz), 2 s in duration, were presented in random order, interspersed with grey screens (4 s duration). Gratings within a trial were randomly assigned one common spatial frequency (0.02, 0.04, 0.16, or 0.32 cpd). For natural stimuli, a 60s movie was presented of an outdoor movie of movement through a field and vegetation. For dark conditions, no visual stimulus was presented (screen off) and all sources of light were switched off.

### Pupil measurements

In some experiments the pupil was monitored using a USB camera (USB 2.0 monochrome camera; ImagingSource) with frames captured at 30 Hz.

### Serum leptin measurements

Serum leptin levels were analyzed using ELISA kits (Mouse Leptin Quantikine Elisa Kit, R&D Systems, USA) according to manufacturer’s instruction. For serum collection, trunk blood was collected from mice that were briefly anesthetized (isoflurane) prior to decapitation. Blood was allowed to clot at room temperature for 1-2 hours and then centrifuged for 20 minutes at 2000 x g. The serum was drawn off and stored at -80°C until used.

### AMPK and PPARα measurements

Levels of total AMPK, phosphorylated AMPK (Thr172), and nuclear PPARα in V1 were measured using ELISA kits according to manufacturer’s instructions (total AMPK: ab181422, Abcam, UK; phosphorylated AMPK: KHO0651, ThermoFisher, UK; PPARα: ab133107, Abcam, UK). For tissue collection, mice were briefly anesthetized (isoflurane) and decapitated. The visual cortex was dissected out, snap frozen using dry ice, and stored at -80°C until used.

### RNA sequencing

Bulk RNA sequencing was performed on V1. V1 tissue was collected and snap frozen from mice that were briefly anesthetized (isoflurane) and decapitated. Tissue was homogenized using a bead mill (Bead Mill 24, Fisher Scientific) and RNA was isolated using an RNeasy kit (Qiagen) according to manufacturer’s instructions. RNA integrity was confirmed (RIN > 7) and mRNA was sequenced at the Oxford Genomics Centre using Illumina NovaSeq 6000.

## QUANTIFICATION AND STATISTICAL ANALYSIS

### Ca^2+^ Imaging Analysis

Image analysis for calcium imaging was performed as previously described^45,47,49^. For motion correct, a discrete Fourier 2D-based image alignment was used (SIMA 1.3.2)^53^. Regions of interest (ROI) were manually drawn around imaged neuronal soma using ImageJ software (NIH public domain; RRID:SCR_002285). Pixel fluorescence was then averaged within each ROI to generate a time series. For each ROI, baseline fluorescence (F_0_) was calculated by taking the 5^th^ percentile of the smoothed time series (1 Hz lowpass, zero-phase, 60^th^-order FIR filter). ΔF/F was then calculated as (F-F_0_/F_0_). FISSA was used to decontaminate the neuropil^55^. Subsequent analyses were performed using custom scripts in MATLAB (MathWorks)^13^.

### Drifting grating analysis

The visual response to drifting gratings was defined as the highest mean ΔF/F, averaged within a 2s window, that occurred within a 4 s window (comprising the 2s grating presentation plus the 2 s of the grey screen presentation which immediately followed the grating presentation). The visual response was subtracted by baseline ΔF/F, defined as the mean value within a 1 s window prior to the visual response.

Responses to gratings of the same angle, but different drift directions were averaged together. Given that multiple spatial frequencies were used, only the spatial frequency with the largest response, meaned across all orientations, was selected for each neuron for subsequent analyses. A neuron’s preferred orientation was that which evoked the largest mean response.

Grating responsive neurons were defined as those for which grating responses were better fit with a double Gaussian curve (direction responses) than with a flat line at zero (null model), which was determined using Bayesian Information Criterion (BIC)^13^. The double Gaussian curve was defined as:

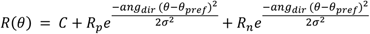

Where R (θ) is the response at a given direction angle θ, C is an offset, R_p_ is the response to the preferred direction after subtracting the offset, θ_pref_ is the angle of the preferred direction, R_n_ is the response to the null direction after subtracting the offset, and *ang_dir_*(*x*) = min (*x*, *x* − 360, *x* + 360), which constrains angular differences to 0-180°, and σ is the standard deviation of the curve.

BIC was given by:

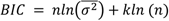

Where n is the number of responses, 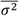 is the mean residual sum of squares of the model, and k is the number of free parameters used by the model, which was zero in the case of the null model. A neuron was considered significantly grating-responsive if the BIC_null_ - BIC_Gaussian_ ≥10, which provides strong evidence against the null model^54^.

### Orientation selectivity

The orientation selectivity index (OSI) of grating-responsive neurons was calculated as:

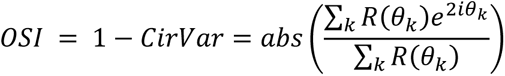

Where *Cirvar* is the circular variance and R (θ_k_) is the mean response to the k_th_ angle θ in orientation space, averaged across direction (see detailed description in ^55^). Mean responses less than zero were set to zero. We used the median OSI values across neurons within an animal.

### Direction selectivity

The direction selectivity index (DSI) of grating-responsive neurons was calculated as:

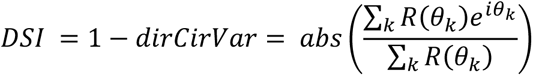

Where *dirCirVar* is the circular variance in direction space and R (θk) is the mean response to the k_th_ angle θ in direction space (see detailed description in ^55^). Mean responses less than zero were set to zero. We used the median DSI values across neurons within an animal.

### Pupil Analysis

Pupil diameter was quantified with custom scripts in MATLAB using functions from the Imaging Processing Toolbox, namely:

1. *imresize* to resize the video
2. *medfilt2* and *imadjust* to remove noise and adjust contrast
3. *imbinarize, imclearborder,* and *bwpropfilt* to find the pupil
4. *regionprops* to fit an ellipsoid around the pupil

The pupil diameter (d) was calculated from the fitted ellipsoid as:

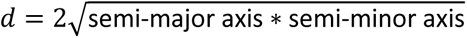

### ATP measurements

For ATeam1.03^YEMK^ imaging, CFP and YFP fluorescence was averaged across ROIs drawn around visible soma in the field of view, after background subtraction. FRET was calculated as a ratio of YFP/CFP fluorescence, and decayed as a function of time with visual stimulation in the presence of ATP synthesis inhibitors. The FRET signal was bound between 0 and 1 by first subtracting the mean FRET signal during the last 3 imaging trials (the FRET signal plateaued at this point), and then dividing by the mean FRET signal at baseline. The FRET decay half time was defined as the time required for the FRET to decay to 50% of its baseline value.

### Differential gene expression analysis

RNA sequencing reads were mapped, using STAR 2.4.oi, to the *Mus musculus* primary assembly (Ensembl v80)^56^. FeatureCounts 1.4.6-p2 was used to count reads uniquely aligned to annotated genes^57^. Differential gene expression analysis was then conducted using DESeq2 1.12.4 (betaPrior = False) to normalize counts^60^. For each gene, a fold change in gene expression was defined as 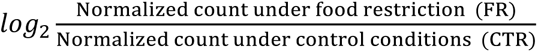 for males and females. Gene set enrichment analysis (GSEA) was conducted using GSEA software (https://www.gsea-msigdb.org/gsea/index.jsp) using the Hallmarks Gene Sets (v7.5.1).

### Statistics

Statistical tests, p values, and the number and definition of independent units are stated in the figure legends. For all measures, the number of animals was taken as the statically independent unit. Sample size was determined by *a priori* based on calculations, with an aim of achieving 80% power on the basis of a 20% group difference and a significance threshold of p < 0.05. Averages are means, except where indicated. Group means and variances were taken from pilot data or previously published data^13^. Where applicable, ANOVA was used to assess significance, followed by *post-hoc* Fisher LSD tests to prevent loss of power. Statistical tests were carried out in Prism 6 (GraphPad Prism; RRID:SCR_002798). Averages denoted in figures represent means, with error bars representing the standard error of the means (S.E.M.). We complemented our statistical analyses using Bayes factor hypothesis testing (Supplemental Figure 3) as described in^35,36^ using JASP software to calculates Bayes factor, effect size, and confidence intervals (https://jasp-stats.org/). As recommended, we used a default, Cauchy distribution with a spread of 1/√2 as a prior distribution of effect size. A BF_10_ < 0.33 is considered to be moderate evidence for the absence of an effect^35,36^.

### Declaration of interests

The authors declare no competing interests.

---

**Supplemental Figure 1.**
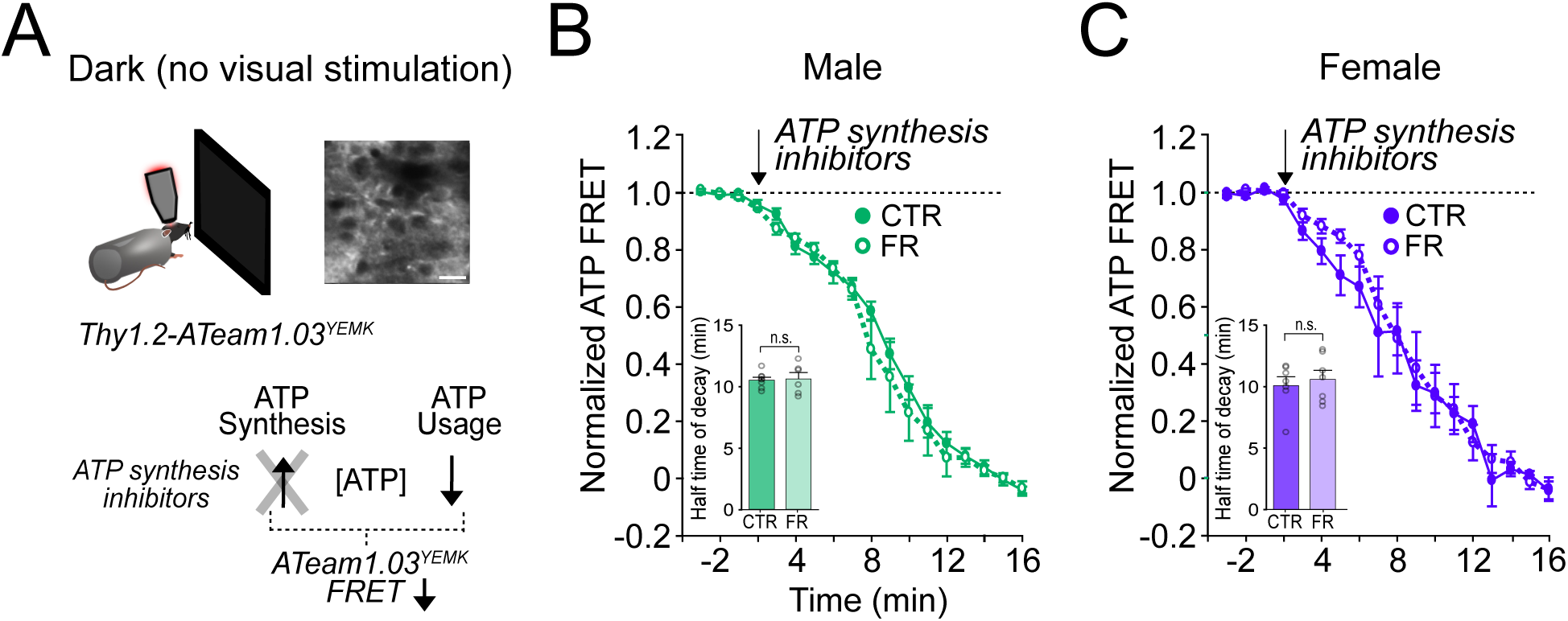
ATP usage in visual cortex is not affected by food restriction in the absence of visual stimulation. (A) Schemata of ATP imaging experiment in the absence of visual stimulation (dark screen). Top: example field of view of V1 layer 2/3 neurons in the ATeam1.03^YEMK^ transgenic mouse (scale bar: 10 μm). Bottom: ATP synthesis inhibitors were used to isolate ATP usage, recorded as a decrease in FRET signal. (B) Normalized ATeam1.03^YEMK^ FRET signal in males in the absence of visual stimulation (dark screen). ATP synthesis inhibitors (arrow) were added to isolate ATP usage. Inset: Time to 50% decay of the ATeam1.03^YEMK^ FRET signal (two-way ANOVA: t = 0.14; df = 24; p = 0. 89; n = 8 CTR males and 6 FR males). (C) As in (B) but for females (inset: two-way ANOVA: t = 0.64; df = 37; p = 0.52; n = 8 CTR females and n = 8 FR females). n.s. = not significant. Error bars are S.E.M.

**Supplemental Figure 2.**
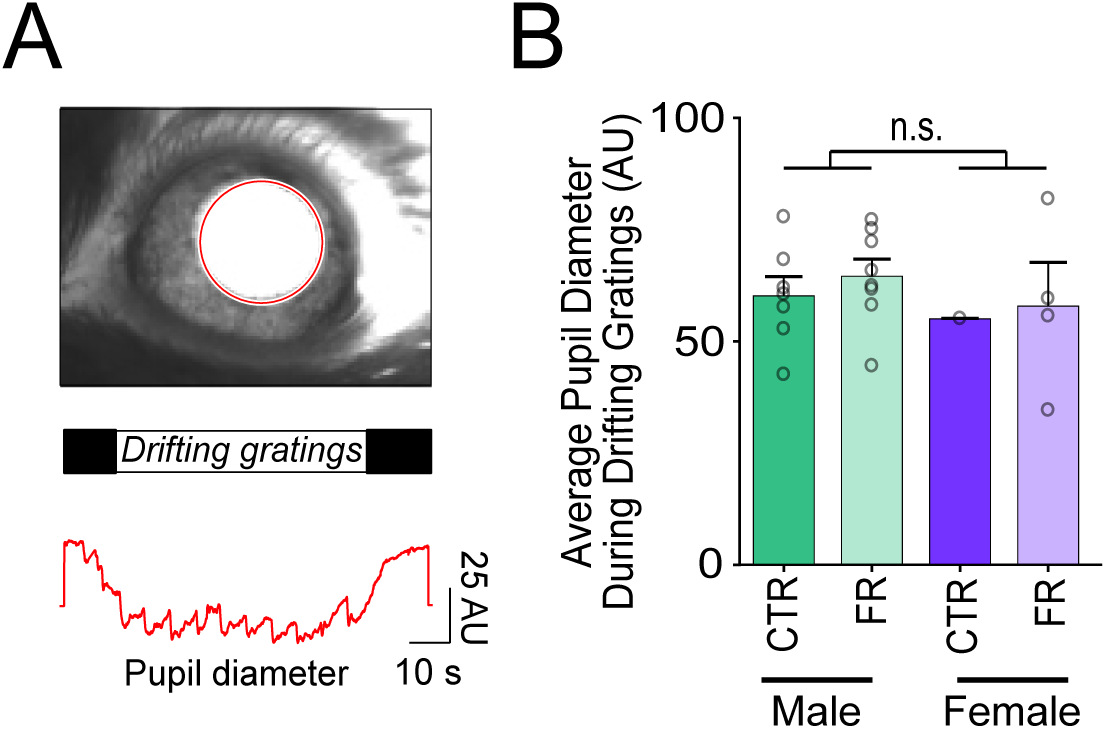
Pupil diameter during visual stimulation is not affected by sex or food restriction. (A) Top: Sample camera image of pupil (red outline) during visual stimulation. Bottom: Pupil diameter during the presentation of drifting gratings. (B) Pupil diameter was comparable across sex and diet. To assess significance across sex, pupil data within a sex was pooled (across diet) (t-test: Male vs Female; t = 0.82; df = 18; p = 0.42; n = 7 CTR males, 8 FR females, 1 CTR female, and 4 FR females). n.s. = not significant. Error bars are S.E.M. Data from male mice are from a previously published dataset obtained under similar conditions^13^.

**Supplemental Figure 3.**
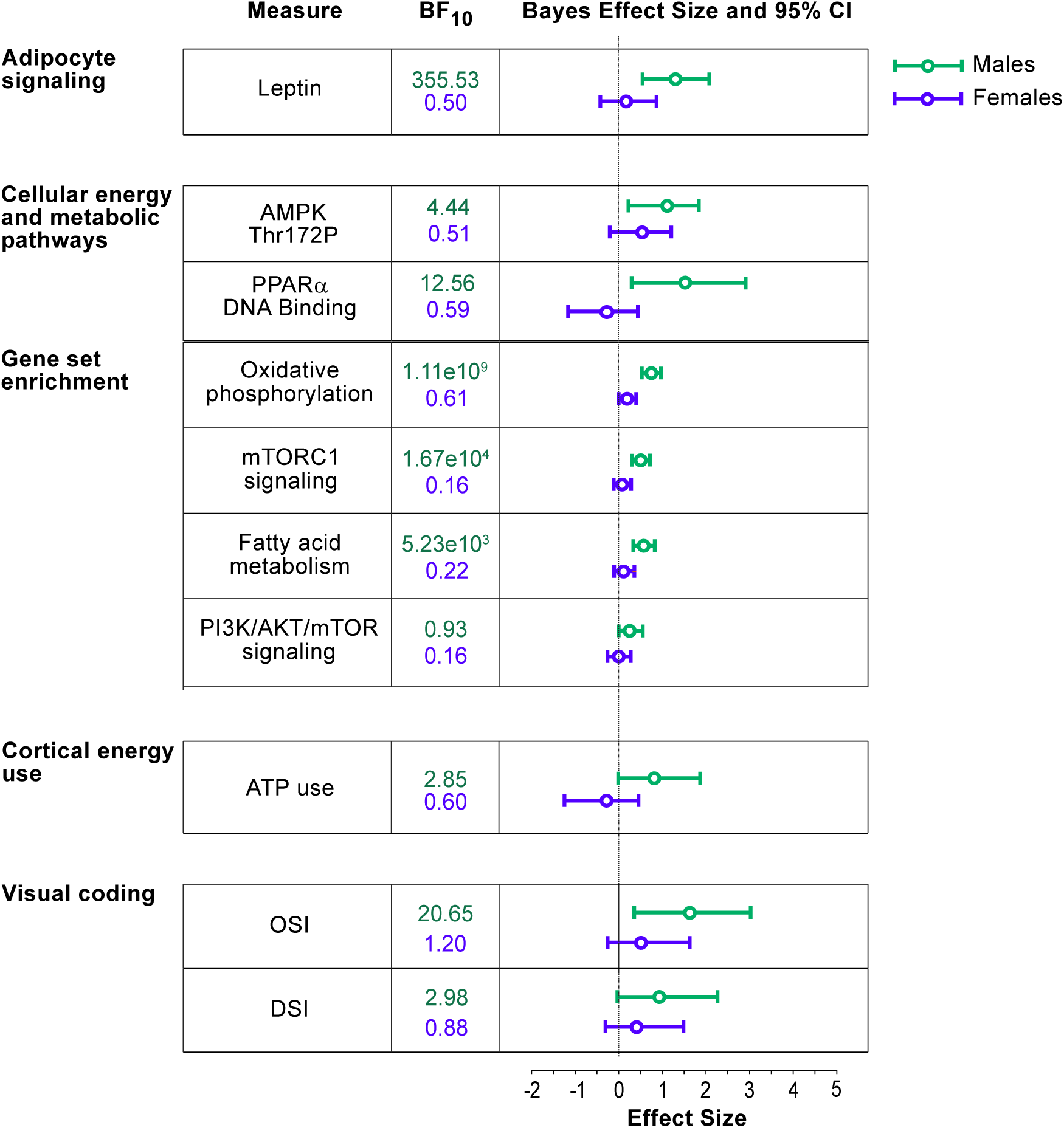
Bayes factor analysis reveals statistically robust impact of metabolic stress on V1 energy usage and visual coding in males, but not in females. Bayes factor (BF_10_) along with quantified effect size and 95% confidence intervals (CI) are shown for males (green) and females (purple). Across parameters, BF_10_ and effect sizes are greater for males than females, meaning that food restriction induces a more robust effect on visual cortical function and energy usage in males than females.

## Notes

### Competing Interest Statement

The authors have declared no competing interest.

### Summary of Updates

We have revised the manuscript adding new experimental results (ATP imaging in Darkness; pupil dilation measurements) and editing the manuscript.

## References

1. Andersson B, Xu XF, Rebuffé-Scrive M, Terning K, Krotkiewski M, Björntorp P. The effects of exercise, training on body composition and metabolism in men and women. Int J Obes. 1991;15(1):75–81.

2. Cortright RN, Koves TR. Sex Differences in Substrate Metabolism and Energy Homeostasis. Can J Appl Physiol. 2000;25(4):288–311.

3. Volek J, Sharman M, Gómez A, et al. Comparison of energy-restricted very low-carbohydrate and low-fat diets on weight loss and body composition in overweight men and women. Nutr Metab (Lond). 2004;1(1):13.

4. McCarthy D, Berg A. Weight Loss Strategies and the Risk of Skeletal Muscle Mass Loss. Nutrients. 2021;13(7).

5. Tirosh A, De Souza RJ, Sacks F, Bray GA, Smith SR, LeBoff MS. Sex differences in the effects of weight loss diets on bone mineral density and body composition: POUNDS LOST trial. J Clin Endocrinol Metab. 2015;100(6):2463–2471.

6. Suchacki KJ, Thomas BJ, Ikushima YM, et al. The effects of caloric restriction on adipose tissue and metabolic health are sex- and age-dependent. Elife. 2023;12.

7. Boutwell K, Rusch P. Some physiological effects associated with chronic caloric restriction. Am J Physiol. 1948;154(3):517–524.

8. Ahlma RS, Prabakaran D, Mantzoros C, et al. Role of leptin in the neuroendocrine response to fasting. Nature. 1996;382(6588):250-252.

9. Gill C, Rissman E. Female Sexual Behavior Is Inhibited by Short- and Long-term Food Restriction. Physiol Behav. 1997;61(3):387–394.

10. Harris JJ, Jolivet R, Attwell D. Synaptic Energy Use and Supply. Neuron. 2012;75(5):762–777.

11. Attwell D, Laughlin SB. An Energy Budget for Signaling in the Grey Matter of the Brain. J Cereb Blood Flow Metab. 2001;21(10):1133–1145.

12. Herculano-Houzel S. Scaling of Brain Metabolism with a Fixed Energy Budget per Neuron: Implications for Neuronal Activity, Plasticity and Evolution. Perc M, ed. PLoS One. 2011;6(3):e17514.

13. Padamsey Z, Katsanevaki D, Dupuy N, Rochefort NL. Neocortex saves energy by reducing coding precision during food scarcity. Neuron. 2022;110(2):280–296.e10.

14. Baile CA, Della-Fera MA, Martin RJ. Regulation of Metabolism and Body Fat Mass by Leptin. Annu Rev Nutr. 2000;20(1):105–127.

15. Guo Z V, Hires SA, Li N, et al. Procedures for behavioral experiments in head-fixed mice. PLoS One. 2014;9(2):e88678.

16. Goltstein PM, Reinert S, Glas A, Bonhoeffer T, Hübener M. Food and water restriction lead to differential learning behaviors in a head-fixed two-choice visual discrimination task for mice. PLoS One. 2018;13(9).

17. Toth LA, Gardiner TW. Food and Water Restriction Protocols: Physiological and Behavioral Considerations. Contemp Top Lab Anim Sci. Published online 2000.

18. Burgess CR, Livneh Y, Ramesh RN, Andermann ML. Gating of visual processing by physiological need. Curr Opin Neurobiol. 2018;49:16–23.

19. Burgess CR, Ramesh RN, Sugden AU, et al. Hunger-Dependent Enhancement of Food Cue Responses in Mouse Postrhinal Cortex and Lateral Article Hunger-Dependent Enhancement of Food Cue Responses in Mouse Postrhinal Cortex and Lateral Amygdala. Neuron. 2016;91(5):1154–1169.

20. Garza-Lombó C, Schroder A, Reyes-Reyes EM, Franco R. mTOR/AMPK signaling in the brain: Cell metabolism, proteostasis and survival. Curr Opin Toxicol. 2018;8:102–110.

21. Dagon Y, Hur E, Zheng B, Wellenstein K, Cantley LC, Kahn BB. P70S6 kinase phosphorylates AMPK on serine 491 to mediate leptin’s effect on food intake. Cell Metab. 2012;16(1):104–112.

22. Dagon Y, Avraham Y, Magen I, Gertler A, Ben-Hur T, Berry EM. Nutritional Status, Cognition, and Survival. J Biol Chem. 2005;280(51):42142–42148.

23. Takei N, Nawa H. mTOR signaling and its roles in normal and abnormal brain development. Front Mol Neurosci. 2014;7(1 APR):1-12.

24. Herzig S, Shaw RJ. AMPK: Guardian of metabolism and mitochondrial homeostasis. Nat Rev Mol Cell Biol. 2018;19(2):121–135.

25. de la Cruz López KG, Toledo Guzmán ME, Sánchez EO, García Carrancá A. mTORC1 as a Regulator of Mitochondrial Functions and a Therapeutic Target in Cancer. Front Oncol. 2019;9(December):1–22.

26. Unger RH, Zhou YT, Orci L. Regulation of fatty acid homeostasis in cells: Novel role of leptin. Proc Natl Acad Sci U S A. 1999;96(5):2327–2332.

27. Poulsen L la C, Siersbæk M, Mandrup S. PPARs: Fatty acid sensors controlling metabolism. Semin Cell Dev Biol. 2012;23(6):631–639.

28. Wójtowicz S, Strosznajder AK, Jeżyna M, Strosznajder JB. The Novel Role of PPAR Alpha in the Brain: Promising Target in Therapy of Alzheimer’s Disease and Other Neurodegenerative Disorders. Neurochem Res. 2020;45(5):972–988.

29. Kwon O, Kim KW, Kim MS. Leptin signalling pathways in hypothalamic neurons. Cell Mol Life Sci. 2016;73(7):1457–1477.

30. Brown LM, Clegg DJ. Central effects of estradiol in the regulation of food intake, body weight, and adiposity. J Steroid Biochem Mol Biol. 2010;122(1-3):65–73.

31. Estep PW, Warner JB, Bulyk ML. Short-Term Calorie Restriction in Male Mice Feminizes Gene Expression and Alters Key Regulators of Conserved Aging Regulatory Pathways. Orban L, ed. PLoS One. 2009;4(4):e5242.

32. Wood SH, Dam S van, Craig T, et al. Transcriptome analysis in calorie-restricted rats implicates epigenetic and post-translational mechanisms in neuroprotection and aging. Genome Biol. 2015;16(1):1–18.

33. Trevisiol A, Saab AS, Winkler U, et al. Monitoring ATP dynamics in electrically active white matter tracts. Elife. 2017;6.

34. Baeza-Lehnert F, Saab AS, Gutiérrez R, et al. Non-Canonical Control of Neuronal Energy Status by the Na + Pump. Cell Metab. 2019;29(3):668–680.e4.

35. van Doorn J, van den Bergh D, Böhm U, et al. The JASP guidelines for conducting and reporting a Bayesian analysis. Psychon Bull Rev. 2021;28(3):813–826.

36. Keysers C, Gazzola V, Wagenmakers EJ. Using Bayes factor hypothesis testing in neuroscience to establish evidence of absence. Nat Neurosci. 2020;23(7):788–799.

37. Collet TH, Sonoyama T, Henning E, et al. A metabolomic signature of acute caloric restriction. J Clin Endocrinol Metab. 2017;102(12):4486–4495.

38. Cuevas-Cervera M, Perez-Montilla J, Gonzalez-Muñoz A, Garcia-Rios M, Navarro-Ledesma S. The Effectiveness of Intermittent Fasting, Time Restricted Feeding, Caloric Restriction, a Ketogenic Diet and the Mediterranean Diet as Part of the Treatment Plan to Improve Health and Chronic Musculoskeletal Pain: A Systematic Review. Int J Environ Res Public Health. 2022;19(11):6698.

39. Hofer SJ, Carmona-Gutierrez D, Mueller MI, Madeo F. The ups and downs of caloric restriction and fasting: from molecular effects to clinical application. EMBO Mol Med. 2022;14(1).

40. Mitchell SJ, Madrigal-Matute J, Scheibye-Knudsen M, et al. Effects of Sex, Strain, and Energy Intake on Hallmarks of Aging in Mice. Cell Metab. 2016;23(6):1093–1112.

41. Kane AE, Sinclair DA, Mitchell JR, Mitchell SJ. Sex differences in the response to dietary restriction in rodents. Curr Opin Physiol. 2018;6:28–34.

42. Gao Q, Horvath TL. Cross-talk between estrogen and leptin signaling in the hypothalamus. Am J Physiol - Endocrinol Metab. 2008;294(5).

43. Irwin RW, Yao J, Hamilton RT, Cadenas E, Brinton RD, Nilsen J. Progesterone and estrogen regulate oxidative metabolism in brain mitochondria. Endocrinology. 2008;149(6):3167–3175.

44. Brinton RD. Estrogen regulation of glucose metabolism and mitochondrial function: Therapeutic implications for prevention of Alzheimer’s disease. Adv Drug Deliv Rev. 2008;60(13-14):1504–1511.

45. Baar EL, Carbajal KA, Ong IM, Lamming DW. Sex- and tissue-specific changes in mTOR signaling with age in C57BL/6J mice. Aging Cell. 2016;15(1):155–166.

46. Brown KD, Waggy ED, Nair S, et al. Sex Differences in Cardiac AMP-Activated Protein Kinase Following Exhaustive Exercise. Sport Med Int Open. 2020;4(01):E13–E18.

47. Clemens AM, Lenschow C, Beed P, et al. Estrus-Cycle Regulation of Cortical Inhibition. Curr Biol. 2019;29(4):605–615.e6.

48. Lin WC, Liu C, Kosillo P, et al. Transient food insecurity during the juvenile-adolescent period affects adult weight, cognitive flexibility, and dopamine neurobiology. Curr Biol. 2022;32(17):3690–3703.e5.

49. de Carvalho T. Calorie restriction or dietary restriction: how far they can protect the brain against neurodegenerative diseases? Neural Regen Res. 2022;17(8):1640.

50. Schroeder JE, Richardson JC, Virley DJ. Dietary manipulation and caloric restriction in the development of mouse models relevant to neurological diseases. Biochim Biophys Acta - Mol Basis Dis. 2010;1802(10):840–846.

51. Pakan JM, Lowe SC, Dylda E, et al. Behavioral-state modulation of inhibition is context-dependent and cell type specific in mouse visual cortex. Elife. 2016;5:1–18.

52. Pakan JMP, Currie SP, Fischer L, Rochefort NL. The Impact of Visual Cues, Reward, and Motor Feedback on the Representation of Behaviorally Relevant Spatial Locations in Primary Visual Cortex. Cell Rep. 2018;24(10):2521–2528.

53. Henschke JU, Dylda E, Katsanevaki D, et al. Reward Association Enhances Stimulus-Specific Representations in Primary Visual Cortex. Curr Biol. 2020;30(10):1866–1880.e5.

54. Kass RE, Raftery AE. Bayes Factors. J Am Stat Assoc. 1995;90(430):773–795.

55. Mazurek M, Kager M, Van Hooser SD. Robust quantification of orientation selectivity and direction selectivity. Front Neural Circuits. 2014;Volume 8, article 92.

56. Dobin A, Davis CA, Schlesinger F, et al. STAR: Ultrafast universal RNA-seq aligner. Bioinformatics. 2013;29(1):15–21.

57. Liao Y, Smyth GK, Shi W. FeatureCounts: An efficient general purpose program for assigning sequence reads to genomic features. Bioinformatics. 2014;30(7):923–930.

